# Conformational dynamics underlying Atypical Chemokine Receptor 3 activation

**DOI:** 10.1101/2023.07.17.549382

**Authors:** Omolade Otun, Christelle Aljamous, Elise Del Nero, Marta Arimont-Segura, Reggie Bosma, Barbara Zarzycka, Tristan Girbau, Cédric Leyrat, Chris de Graaf, Rob Leurs, Thierry Durroux, Sebastien Granier, Xiaojing Cong, Cherine Bechara

**Author notes:** Sosei Heptares, Steinmetz Building, Granta Park, Great Abington, Cambridge, CB21 6DG, UK.

## Abstract

Atypical Chemokine Receptor 3 (ACKR3) is a G protein-coupled receptor that does not signal through G proteins. It is known as a chemokine scavenger involved in various pathologies, making it an appealing yet intriguing therapeutic target. Indeed, the structural properties that govern ACKR3 functional selectivity and the overall conformational dynamics of ACKR3 activation are poorly understood. Here we combined Hydrogen/Deuterium exchange mass spectrometry (HDX-MS) and molecular dynamics simulations to examine the binding mode and mechanism of action of various small-molecule ACKR3 ligands of different efficacy for β-arrestin recruitment. Our results show that activation or inhibition of ACKR3 is largely governed by intracellular conformational changes of helix 6, intracellular loop 2 and helix 7, while the DRY motif becomes protected during both processes. Moreover, HDX-MS identifies the binding sites and the allosteric modulation of ACKR3 upon β-arrestin 1 binding. In summary, this study highlights the structure-function relationship of small-molecule ligands, the overall activation dynamics of ACKR3, the binding mode of β-arrestin 1 and the atypical dynamic features in ACKR3 that may contribute to its inability to activate G proteins.

## Introduction

Chemokine receptors are class A G protein-coupled receptors (GPCRs) that are critical for immune cell migration, immune modulation, wound healing, inflammation and host pathogen interactions^1^. Most chemokine receptors are able to signal through the canonical G-protein mediated pathways. However, a small subset of ‘atypical’ chemokine receptors (ACKRs) is unable to activate G-proteins. ACKRs shape chemokine gradients via chemokine scavenging and dampen inflammation through β-arrestin-dependent internalization pathways^2^. Among these, ACKR3 (formerly known as CXCR7)^3^ has also been reported to bind non-chemokine ligands and scavenge opioid peptides^4^. ACKR3 also interact with various other membrane receptors (e.g. CXCR4) and alter their subcellular distribution and signalling^4^. Despite possessing the general architecture and conserved sequence motifs of class A GPCRs, ACKR3 appears to exclusively activate β-arrestins^5, 6^. Several cryo-EM structures of ACKR3 were recently resolved in active state, bound to its chemokine ligand CXCL12 and/or a small-molecule agonist CCX662^7^. These structures exhibit the hallmarks of canonical class A GPCR activation, mainly the outward displacement of helix 6 (H6) on the intracellular side. Yet, several distinct features were observed, including the atypical orientation of CXCL12, a short helix in the intracellular loop 3 (ICL3) - or rather a kink at the C-terminus of H6 - and the lack of a kink at the N-terminus of H4 into ICL2. It is unclear whether these features are responsible for the atypical function of ACKR3 or are rather associated with the binding of the synthetic Fabs. In the cryo-EM structure without the intracellular Fabs, ICL1-3 and H8 all appeared disordered. A recent NMR study^8^ highlighted that agonist-bound ACKR3 conformations differed from the antagonist-bound ones at the intracellular probe M138^3.46^ (superscript refers to Ballesteros and Weinstein nomenclature^9^), a known microswitch of class A GPCR activation. This reflected the H6-H7 movements upon ACKR3 activation, as later revealed by the cryo-EM structures. Kleist *et al.* also found that point mutations at N127^3.35^, part of the sodium-binding site, drastically affected ACKR3 activation, similar to previous findings in CXCR4^10^. Therefore, it is still unclear which structural properties distinguish ACKR3 from canonical class A GPCRs and confer its selectivity towards β-arrestin signalling. Moreover, structural features of the inactive ACKR3 state are unknown such that the molecular bases underlying the transition between inactive and active conformations are poorly understood and require further investigation.

GPCRs activation is a dynamic process during which the receptor oscillates between discrete conformational states, the most populated free energy state being in general the lowest-energy inactive state^11, 12^. Ligand binding induces conformational changes in the receptor, shifting its conformational equilibrium and influencing its activity depending on the ligand efficacy. Advances in structural biology approaches have provided great insights into the diversity of GPCRs conformations^13, 14^. Numerous high resolution GPCR structures of various conformational states have revealed high diversities in ligand recognition yet striking similarities in receptor conformational changes upon activation. However, cryo-EM and X-ray crystal structures are static snapshots of the most stable and/or accessible states under the experimental conditions used. Therefore, to understand the full complexity of GPCR conformational landscapes, complementary biophysical techniques are required to address transient states that might be discarded during sample preparation or data processing stages^15–22^. Importantly, integration of several of these biophysical and computational techniques have proven especially useful for probing activation dynamics of several GPCRs^23–29^.

Hydrogen/Deuterium exchange coupled with mass spectrometry (HDX-MS) is one such approach that provides a comprehensive dynamic view of GPCR structural transitions upon binding to various ligands^30–35^ or intracellular partners^36, 37^. HDX-MS probes the exchange between backbone amide hydrogens and deuterium present in the solvent, which is mainly related to the amide hydrogen solvent accessibility and hydrogen bond (H-bond) stability.

Therefore, it is a label-free method that reflects the stability of protein secondary structures, the binding interface with partners, as well as the overall protein conformational dynamics^38, 39^. When coupled with molecular dynamics (MD) simulations^40^, HDX-MS offers the validation of the MD predictions, which together can reveal detailed receptor-ligand interactions and the conformational dynamics underlying receptor activation and inhibition.

Here, we combine HDX-MS analysis and MD simulations to study the conformational changes of ACKR3 induced by a small-molecule agonist and two inverse agonists. These changes revealed important structural features associated with ACKR3 activation, including both commonalities with class A GPCRs and unique features specific to ACKR3. Namely, the agonist altered the receptor conformation at the orthosteric pocket, triggering H6 opening on the intracellular side and high mobility of ICL2, which corresponds to increased deuterium uptake in the latter two regions. Inverse agonists, by contrast, locked the receptor in an inactive state and decreased the deuterium uptake throughout the receptor. In addition, all ligands resulted in shielding of the DRY motif. β-arrestin 1 binding significantly reduced the deuterium uptake of the intracellular loops of ACKR3, suggesting therefore a core engagement. The NPxxY and the DRY motifs, on the other hand, became more deuterated in the presence of β-arrestin 1, implying that these motifs are not directly implicated in the binding to β-arrestin 1. Overall, our results provide insights into the activation mechanism of ACKR3 and its propensity towards β-arrestin recruitment.

## Results

### Optimization of HDX-MS to probe ACKR3 conformational changes upon ligand binding

We developed an HDX-MS strategy using detergent-purified ACKR3 to identify the dynamic conformational changes in ACKR3 upon ligand binding. Three small-molecule ligands were studied, which had been characterised for their pharmacological effect towards β-arrestin recruitment: an agonist VUF15485^41, 42^ and two inverse agonists VUF16840 and VUF25550^43–45^ (Fig. 1A). All tested ligands have been shown to compete with CXCL12, an endogenous ACKR3 ligand. Extensive optimization of the receptor purification and digestion conditions allowed identification of 100 to 140 peptides covering around 90% of the receptor sequence, with an average peptide redundancy above three, after manual inspection (HDX summary tables, Supplementary data 1).

**Figure 1.**
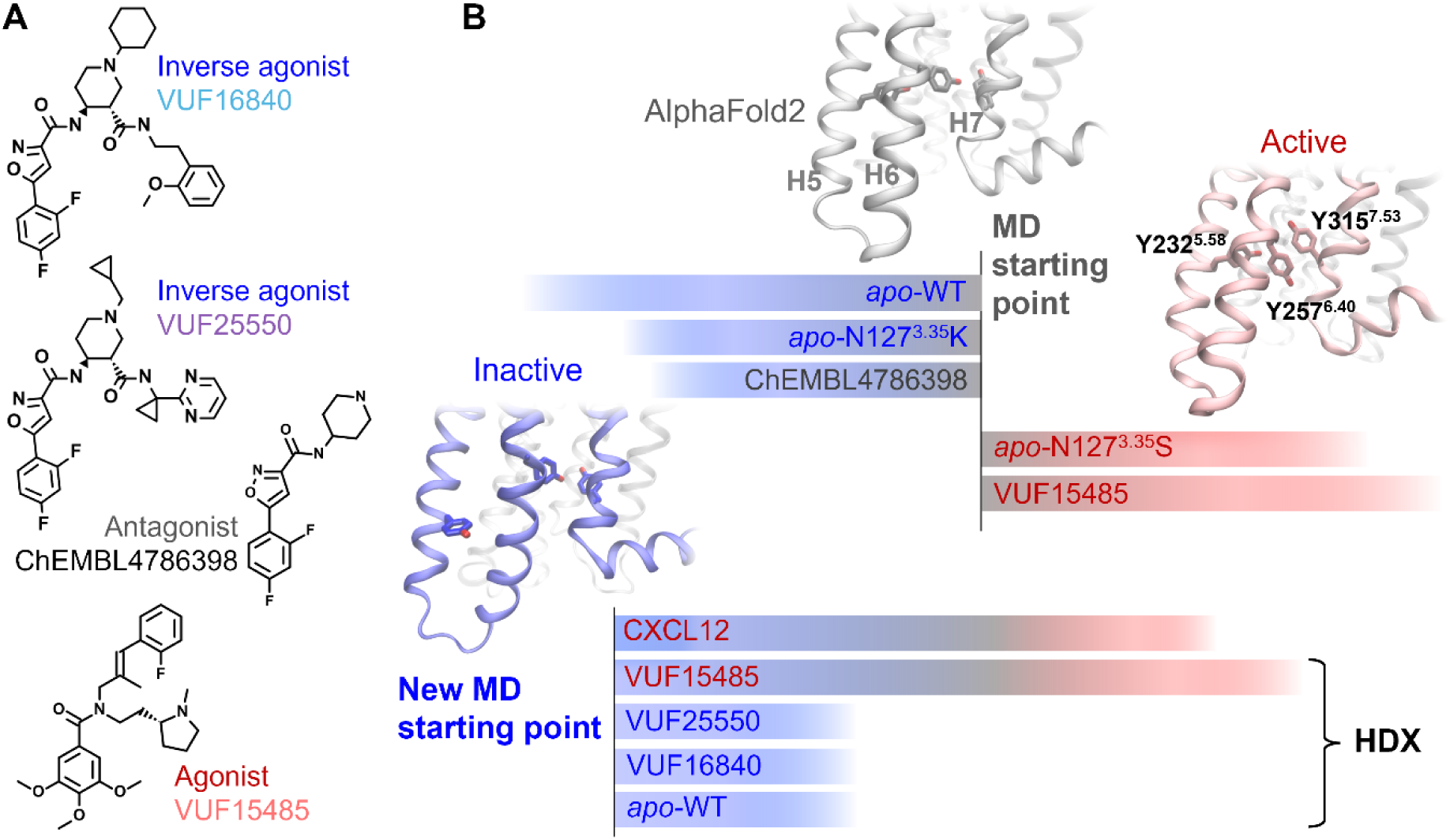
Small-molecule ligands and MD systems studied. **A)** Chemical structures of the agonist VUF15485, inverse agonists VUF16840 and VUF25550, and an antagonist ChEMBL4786398. ChEMBL4786398 is a substructure of the inverse agonists, which was used here to facilitate the binding mode prediction. **B)** Two clusters of MD simulations from different starting structures. Initially, five simulations were performed starting from the AF model, including WT ACKR3, a constitutively active mutant (N127^3.35^S), an inactive mutant (N127^3.35^K) in *apo* form, as well as WT ACKR3 bound to VUF15485 and ChEMBL4786398. The agonist-bound receptor and the constitutively active mutant evolved toward active states. The other three systems evolved toward inactive states featured by H6 inward movements. The final inactive state of *apo* WT ACKR3 was then used as the new starting point to investigate the receptor conformational changes induced by agonists and inverse agonists. HDX protection factors were calculated from the new trajectories using the *apo* form as a reference state. These were compared with the HDX-MS data to validate the MD sampling.

In order to determine the effects of ligand binding on the ACKR3 conformations, we performed differential HDX (ΔHDX) analysis between *apo* and ligand-bound ACKR3. Two to three biological replicates were performed for every condition (HDX summary tables and data tables, Supplementary data 1 and 2). Overall, the receptor N-terminus, C-terminus and loops showed higher overall deuteration due to their unstructured properties and high flexibility (Fig. S1). Conversely, we observed lower overall deuteration in the peptides corresponding to the helical transmembrane domains, reflecting their well-structured nature and low solvent accessibility due to the presence of the detergent micelle (Fig. S1). Changes in these regions upon adding the ligands, if present, were thus mostly observed at longer deuteration times. We next used the overall ΔHDX experimental data to validate the MD simulations of ACKR3 in the presence of the three different ligands.

### ACKR3 in inactive state conformation

MD simulations were performed to study the conformations of ACKR3 in *apo* and ligand-bound forms. We used the REST2 technique^46^ to enhance the MD sampling (see Methods for details). HDX protection factors (logPF) were calculated from the simulation trajectories using HDXer^47^, which enabled a direct qualitative comparison with the deuterium uptake data from HDX-MS. This validated the MD prediction of the ligand binding poses and receptor conformational changes, and gave insights into the mechanism of action of the agonist *versus* inverse agonists.

The receptor conformation in *apo* form is a key reference for the calculation of ligand-induced HDX protection factors. We first attempted to obtain the *apo* conformations by REST2 MD simulations starting from the active state in the cryo-EM structures of ACKR3^7^. However, due to the short helix at ICL3 (or H6 kink), the receptor showed high mobility in this region and unfolding of H6 even in the presence of inverse agonists. This suggests that the H6 kink is unfavourable without the intracellular Fab used for the cryo-EM structure. We therefore built an AlphaFold2 (AF) model of ACKR3, which turned out to be nearly identical to the cryo-EM structures except for the H6-ICL3 region. It has a longer H6 until residue K247^6.30^ and a shorter ICL3 without the helical kink, similar to that in the inactive structures of CXCR1, CXCR2 and CXCR4 (PDBs 2LNL, 6LFL and 3ODU). Using the AF model as a starting point of MD simulations, we found that the *apo* receptor stabilized into an inactive conformation typical of class A GPCRs, in which H6 moved significantly inward on the intracellular side (Figs. 1 and S2). To assess that the H6-closed conformation indeed represented the inactive state, we introduced various ligands and point mutations into the AF model and performed the same simulations. The antagonist and inactive mutation also led to the H6 inward movements (Figs. 1 and S2). By contrast, the agonist and the constitutively active mutation resulted in further openings of H6 on the intracellular side. The agonist-bound ACKR3 also exhibited Y232^5.58^-Y257^6.40^-Y315^7.53^ π-stacking as H6 opened, same as in the CXCL12-bound cryo-EM structure (Fig. 1 and S2). These results suggest that ACKR3 conserves not only the sequence motifs of class A GPCRs but also the common conformational changes upon activation, whereas the initial AF model may represent an intermediate or “active-like” state. The *apo* ACKR3 model in inactive state then served as the reference *apo* state and a new starting point of MD simulations, to study the impact of different ligands on the receptor conformation. The MD simulations of ACKR3 bound to ChEMBL4786398 served to deduce the binding mode of the more challenging analogs, VUF16840 and VUF25550 (Fig. 1A). The MD-predicted ligand binding mode and receptor dynamics were in good agreement with the HDX-MS and site-directed mutagenesis data, as discussed below.

### Ligand binding mode and protection at ACKR3 orthosteric site

The HDX-MS data showed a decreased deuterium uptake at the extracellular face of ACKR3 in the presence of all tested ligands, regardless of their pharmacological profile (Figs. 2 and S3). Namely, we observed protection to deuteration at the level of the receptor orthosteric binding pocket. The most significant protection was at several peptides spanning the extracellular top of H5. In the case of peptide 214-220 at H5, protection was up to 35% and 24% by the two inverse agonists and 17% by the agonist. The top of H4 was also highly protected, mainly with the inverse agonists (Fig. 2). For peptide 175-181 at H4 for instance, inverse agonists VUF25550 and VUF16840 induced a decrease in deuteration whereas the agonist VUF15485 induced a slight protection only at longer deuteration times.

**Figure 2.**
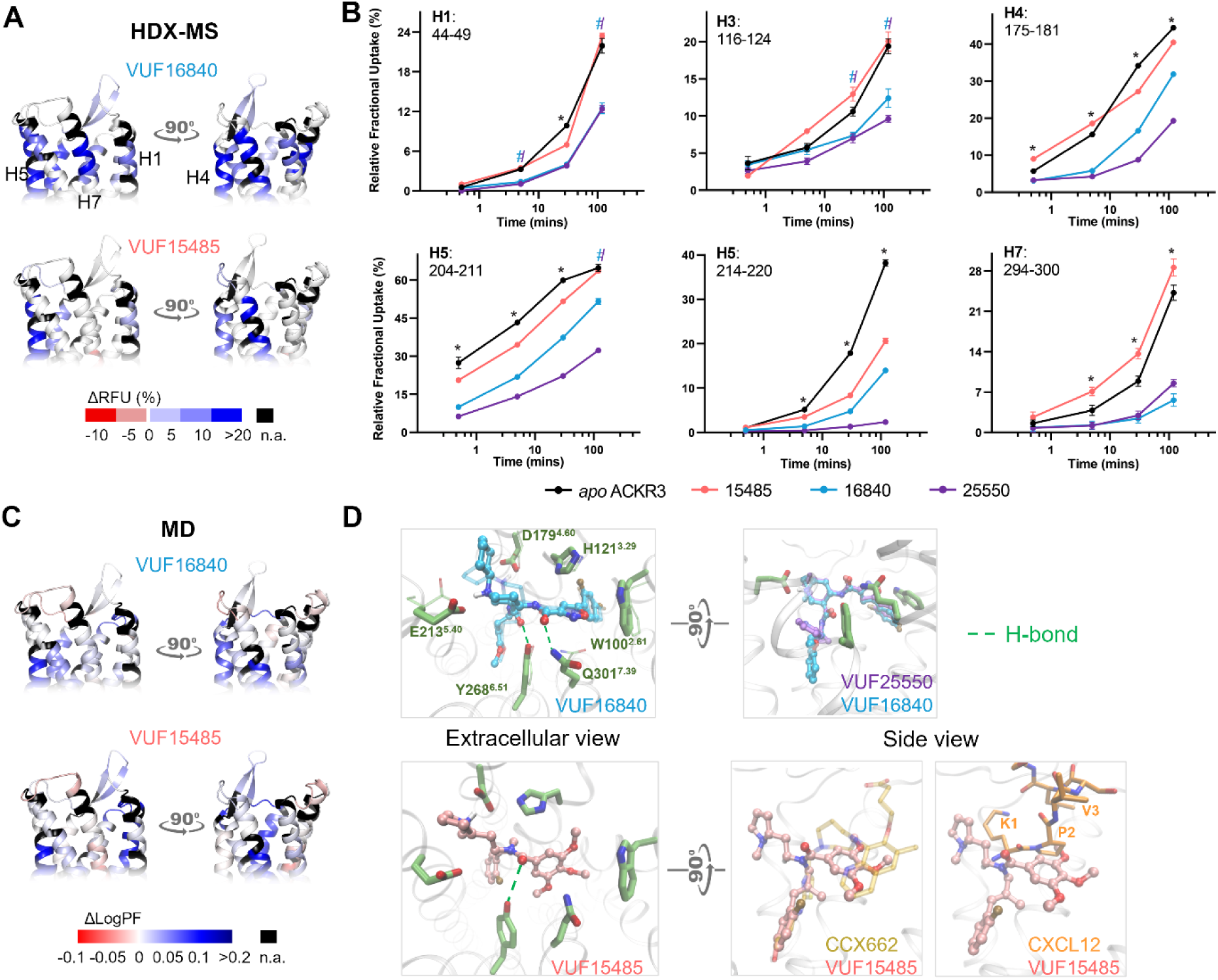
Ligand binding at the level of ACKR3 orthosteric pocket. **A)** Schematic representation of the % differential relative fractional uptake data (apo ACKR3 – bound ACKR3) mapped onto the upper part of the AF model of ACKR3 (for clarity, the N-terminus is not shown). This depicts reproducible and statistically significant ΔHDX in response to inverse agonists small ligands represented by VUF16840 (VUF25550 giving a similar profile), or agonist small ligand VUF15485. Black regions represent regions with no sequence coverage. Ligand-induced reduction in deuterium uptake is represented in blue whilst ligand-induced increase in deuterium uptake is in red, according to the scale. **B)** Associated deuterium uptake plots showing the relative uptake for peptides from apo or ligand-bound ACKR3 across several deuteration time points and are representative of the extracellular region as indicated at the top of the plot. Statistically significant changes were determined using Deuteros 2.0 software^48^ (p ≤ 0.01): black stars represent changes significant for all 3 ligands, and two barred crosses with the related ligand colour represent changes significant for one or two ligands. Uptake plots are the average and SD of 3 technical replicates from the same biological preparation of ACKR3. **C)** Calculated differential log HDX protection factor changes (ΔlogPF) mapped on the AF model. Per-residue logPF was first calculated for each MD trajectory of ACKR3 in *apo* and ligand-bound forms. The difference between the *apo* and bound forms gave the per-residue ΔlogPF for each ligand. For comparison with the HDX data, per-peptide ΔlogPF was calculated by averaging the per-residue ΔlogPF over the peptides obtained in the HDX-MS experiments for each ligand. **D)** Predicted ligand binding mode. The inverse agonists could bind in both enantiomers of the piperidine group, illustrated here with VUF16840. VUF25550 bound similarly to VUF16840. Their difluorophenyl formed π–π stacking with W100^2.61^ and more contacts with H1, H2 and H7 than the agonist. The agonist VUF15485 formed ionic interactions with D179^4.60^ via its 1-methylpyrrolidine. The rest of VUF15485 largely overlaps with CCX662 as shown in the superimposition to PDB 7SK8. It also overlaps with the N-terminus of CXCL12.

The MD prediction of ligand binding mode and HDX protection factors were coherent with the HDX-MS data (Fig. 2C). All the 3 small-molecule ligands anchored to D179^4.60^ via the amine (Fig. 2D), resembling the role of residue K1 in CXCL12^7^. The inverse agonists, VUF16840 and VUF25550, could bind in both enantiomers of the amine, which anchored to D179^4.60^ and E213^5.40^ respectively. The agonist VUF15485, however, was only stable in one enantiomer in the simulations, which anchored to D179^4.60^. VUF15485 occupied the pocket space that largely overlaps with the partial agonist CCX662 and the N-terminus of CXCL12 in the cryo-EM structures (PDBs 7SK8 and 7SK3). It formed a H-bond with Y268^6.51^ via its amide carbonyl. The two inverse agonists extended further in the pocket toward H1, H2 and H7. As close analogs, they exhibited nearly identical binding poses (Fig. 3A), each forming two H-bonds with Y268^6.51^ and Q301^7.39^. The predicted ligand binding poses are confirmed by site-directed mutagenesis data on VUF15485 and VUF16840 binding (Table S1). Namely, a D179^4.60^N mutation diminished the binding affinities for both ligands, whereas E213^5.40^Q only affected VUF16840. In addition, Q301^7.39^E/A mutations diminished the affinity for VUF16840 but not VUF15485. A D275^6.58^N mutation had no impact on either ligand, since the mutation site lies on the extracellular rim of the orthosteric pocket, distant from the predicted ligand-binding site. Several other mutations in the orthosteric pocket were tested to validate the binding pose of VUF15485^42^.

**Figure 3.**
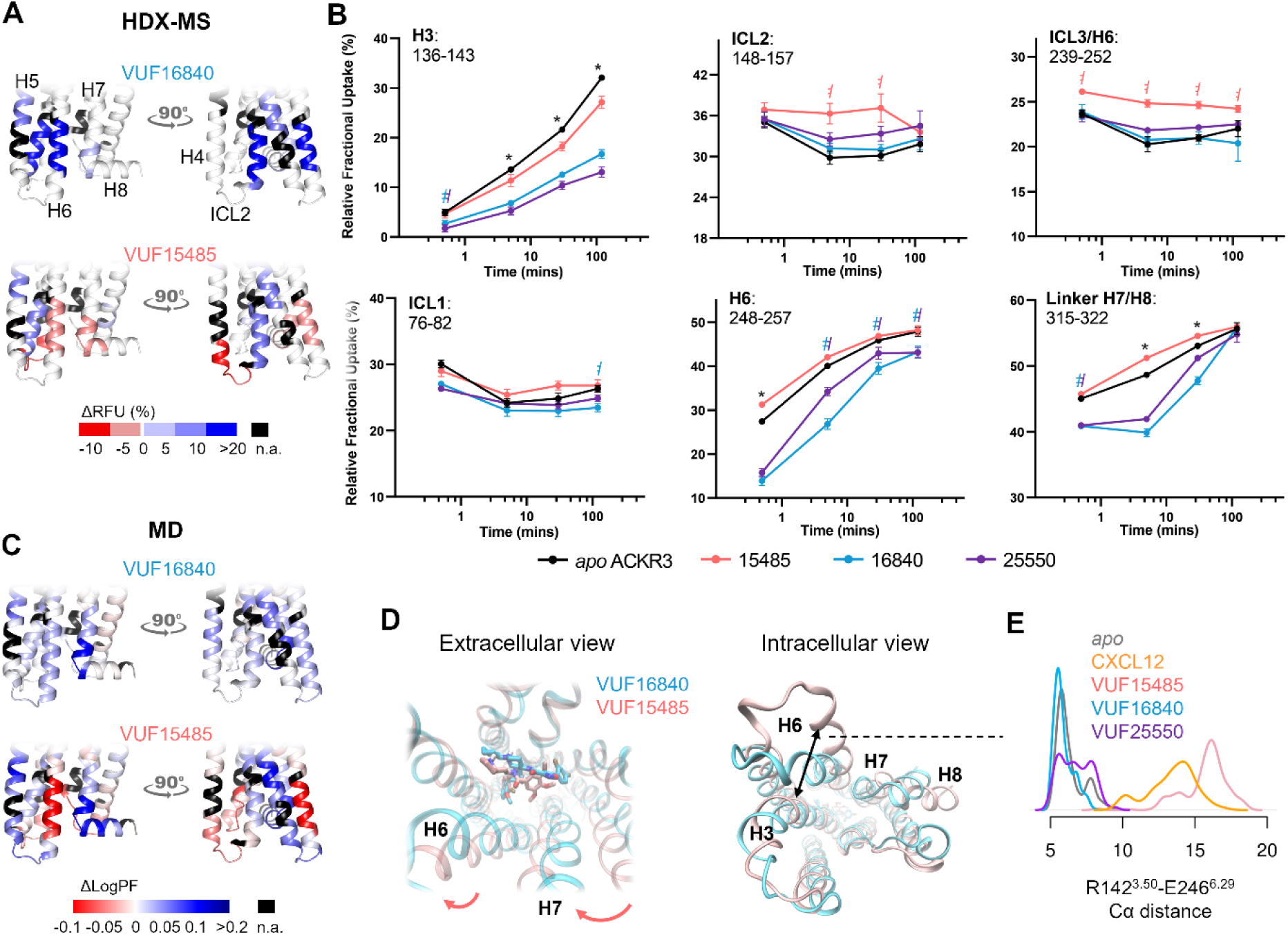
Allosteric conformational changes of ACKR3 activation. **A)** Schematic representation of the % differential relative fractional uptake data (apo ACKR3 – bound ACKR3) mapped onto the lower part of the AF model of ACKR3 (for clarity, the C-terminus is not shown). This depicts reproducible and statistically significant ΔHDX in response to inverse agonists small ligands represented by VUF16840 (VUF25550 giving a similar profile), or agonist small ligand VUF15485. Black regions represent regions with no sequence coverage. Ligand-induced reduction in deuterium uptake is represented in blue whilst ligand-induced increase in deuterium uptake is in red, according to the scale. **B)** Associated deuterium uptake plots showing the relative uptake for peptides from apo or ligand-bound ACKR3 across several deuteration time points and are representative of the intracellular region as indicated at the top of the plot. Statistically significant changes were determined using Deuteros 2.0 software^48^ (p ≤ 0.01): black stars represent changes significant for all 3 ligands, and two barred crosses with the related ligand colour represent changes significant for one or two ligands. Uptake plots are the average and SD of 3 technical replicates from the same biological preparation of ACKR3. **C)** Calculated ΔlogPF from the MD simulations, mapped on the AF model. **D)** Proposed mechanism of activation. The bulky trimethoxybenzamide group of VUF15485 induces a “twist” of the 7TM bundle around the orthosteric pocket, which allosterically triggers H6 opening on the intracellular side. **E)** Probability density distribution of the ionic-lock residue distances during the MD simulations.

The HDX-MS data at the orthosteric pocket of ACKR3 and the MD predicted ligand binding mode are coherent with the ability for all tested ligands to inhibit CXCL12 binding. Cryo-EM structures of ACKR3 in complex with CXCL12 shows that CXCL12-K1 and P2, as part of the receptor activation motif of CXCL12, penetrate ACKR3 orthosteric pocket to make side chain interactions with E213^5.38^, D179^4.60^ and Y200^ECL2^. CXCL12-V3 backbone amide was also shown to potentially form a H-bond with ACKR3-Q301^7.39^ in addition to packing of V3 side chains against proximal residues, L297^7.35^ and H298^7.36^ (Fig. 2D). MD predictions show that the small-molecule ligands overlap largely with CXCL12-K1 and P2. In correlation, HDX uptake is reduced for peptides spanning these residues where all the ligands show main contacts at the level of H4 and H5. Protective effects on H7 were mainly visible for the inverse agonists which form two additional H-bonds with Q301^7.39^ (Fig. 2). Taken together, our data indicate overlapping contacts for CXCL12 and the small-molecule ligands at the orthosteric binding pocket.

### All ligands shield the DRY motif

ΔHDX-MS showed that binding of the different ligands led to significant allosteric changes in the intracellular side of ACKR3. Notably, the DRY motif at the intracellular face of H3 was protected in the presence of all ligands, implying that this motif is always shielded upon binding of these ligands (Fig. 3). The agonist-induced protection on the representative peptide 138-143 spanning the DRY motif was weaker than the inverse agonists (Figs. 3 and S4). MD simulations suggest that in agonist-bound forms, the DRY is shielded by H5 and Y257^6.40^ (Fig. S4). The Y257^6.40^ orientation here is similar to the one observed in the CXCL12-bound cryo-EM structure. This agrees with the changes in the NMR spectra of M138^3.46^ upon agonist binding, where an aromatic residue was detected in its proximity^8^. Indeed, we observed in the MD simulations that Y257^6.40^ was dynamic in the *apo* form, whereas agonists led to proximation of Y257^6.40^ to M138^3.46^ (Fig. S4A). However, we did not observe a clear correlation between the M138^3.46^ side-chain conformations and the ligand efficacies (Fig. S4B). In the inverse agonist-bound forms, the DRY is shielded by H6 and protected by the overall decrease of receptor conformational dynamics. The DRY motif is highly conserved across GPCRs and has been shown to be critical for both G protein activation and in maintenance of the ionic lock through the hydrophobic network involving D141^3.49^, R142^3.50^ and conserved residues at position 6.30. Particularly, R142^3.50^ has been shown to have direct G protein contacts in structures of several GPCR ternary complexes^49–51^. The current accepted mechanism of G-protein activation by class A GPCRs involves the release of R142^3.50^ from the ionic lock allowing critical interactions required for G protein activation^52^. Since all the ligands we tested induced a protection at this region, this protection could be therefore related to the inability for ACKR3 to activate G proteins.

### Typical and distinct features of ACKR3 activation

The MD simulations of ACKR3 bound to CXCL12 and VUF15485 both displayed a remarkable H6 opening on the intracellular side (Fig. 3D), typical of class A GPCR activation^53^. The intracellular half of H6 was deprotected, as shown by the lower logPF than the *apo* form calculated from the MD trajectories (Fig. 3C). This was confirmed by the HDX-MS results, in which the lower half of H6 showed higher uptake in the presence of VUF15485 compared to the *apo* receptor (Figs. 3A and 3B). Interestingly, ΔHDX showed that I254^6.37^ was protected with inverse agonists and deprotected with the agonist, which correlates with the observed interactions between I254^6.37^ and M138^3.46^ only in inactive structures of GPCRs^8^. The H7-H8 linker, however, showed discrepancies between the MD and the HDX-MS data. The H6 outward and H7 inward movements appear to be overestimated in the agonist-bound simulations compared to the *apo* state. This may be due to the C-terminal truncation in the MD and/or to insufficient sampling of the *apo* state. Although ACKR3 has a high basal activity, the *apo* state did not show activation during the MD simulations, which likely overestimated the ΔlogPF of the agonist-bound state. ICL2 and the intracellular tip of H4 also exhibited significant deprotection upon agonist binding, in both the MD simulations and the HDX-MS experiments (Fig. 3). MD simulations revealed that VUF15485 reduced the H2-H4 and H3-H6 contacts on the intracellular side allosterically (Fig. 3D). As H6 moved outward, the contacts between H5-ICL3-H6 and H3 changed remarkably, leading to ICL2 unfolding. Therefore, ICL2 and the intracellular tip of H4 were more mobile and deprotected. In the CXCL12-bound form, ICL2 also unfolded as H6 moved outward. In *apo* ACKR3 and the inverse agonist-bound forms, ICL2 maintained the initial fold like in the cryo-EM structures bound to an intracellular Fab^7^. Interestingly, ICL2 in most class A GPCRs forms a short α-helix in both inactive and active states. ICL2 has been associated with G protein subtype selectivity and G protein signalling of several receptors^36, 37, 54^. We speculate that the unfolding of ACKR3 ICL2 upon agonist binding may contribute to the lack of G protein signalling of this atypical receptor.

Taken together, our data suggest the following mechanism of activation by the agonist: VUF15485 may induce an overall twist of the 7TM bundle, due to the bulky trimethoxybenzamide ring in the centre of the orthosteric pocket (Fig. 3D). The twist propagates to the intracellular side allosterically in a loosely-coupled manner, which triggers H6 outward movements. A similar phenomenon was observed in the simulations of CXCL12-bound ACKR3 (Fig. S5). The inverse agonists, by contrast, may stabilise the inactive state of the 7TM bundle through additional interactions with the receptor within the orthosteric pocket. Namely, the inverse agonists are less bulky and extend toward H1-H2. They form additional interactions with H2 (π-stacking with W100^2.61^) and H7 (H-bond with Q301^7.39^), which may restrain receptor conformational changes (Fig. 2D). Consistently, the inverse agonists resulted in additional protection on H2 and H7 in the orthosteric pocket, as well as overall protection of the receptor from HDX (Fig. 2). Mutation Q301^7.39^A has been shown to significantly increase ACKR3 basal activity^7^. Here, our findings indicate that Q301^7.39^ is also key for ligand-dependent activity. Indeed, ligand-H7 contacts have been reported to be important for β-arrestin signalling bias in other class A GPCRs^8, 28, 55–57^. Although there appears to be no common pattern across different GPCR families, the pocket area between H2 and H7 is likely a hot spot for ligand bias. The above data show that agonist-induced ACKR3 activation increases the solvent accessibility of H6, ICL2, H7 and the H7-H8 linker on the intracellular side. This suggests that conformational changes occur to facilitate the recruitment of β-arrestin through core engagement.

### HDX-MS identifies putative β-arrestin binding sites

To follow-up on ACKR3 binding to β-arrestin 1, we took advantage of the constitutively active properties of the receptor and performed ΔHDX analysis of ACKR3 in the presence of 1.2 molar equivalent β-arrestin 1 ΔCter (Fig. 4). As the C-tail of β-arrestin has an autoinhibitory function, its removal was necessary to shift β-arrestin 1 towards the active state^58^. The presence of β-arrestin 1 led to the stable protection of ICL2 and ICL1 and a slight protection of ICL3 (Fig. 4). Interestingly, peptides spanning the NPxxY motif as well as the intracellular face of H6 were deprotected. Another notable effect was the deprotection of a representative peptide 90-95 within H2 at the orthosteric pocket. This region is right above the highly conserved D90^2.50^, which is part of the sodium binding site in class A GPCRs. Sodium ions are known negative allosteric modulators of class A GPCR activity^59^. The deprotection observed at this level implies allosteric cooperation between the ligand/sodium-binding sites and β-arrestin binding, which may be explored for ligand design. Another notable finding was the deprotection of the DRY motif in the presence of β-arrestin 1 (representative peptide 138-143, Fig. 4). Therefore, the ACKR3 DRY motif becomes more exposed and/or dynamic upon binding to β-arrestin 1, suggesting that this motif is not implicated in the direct interaction with β-arrestin 1. Finally, we observed some allosteric effects upon ACKR3 binding to β-arrestin 1, the most notable one being the ECL2 protection (Fig. 4).

**Figure 4.**
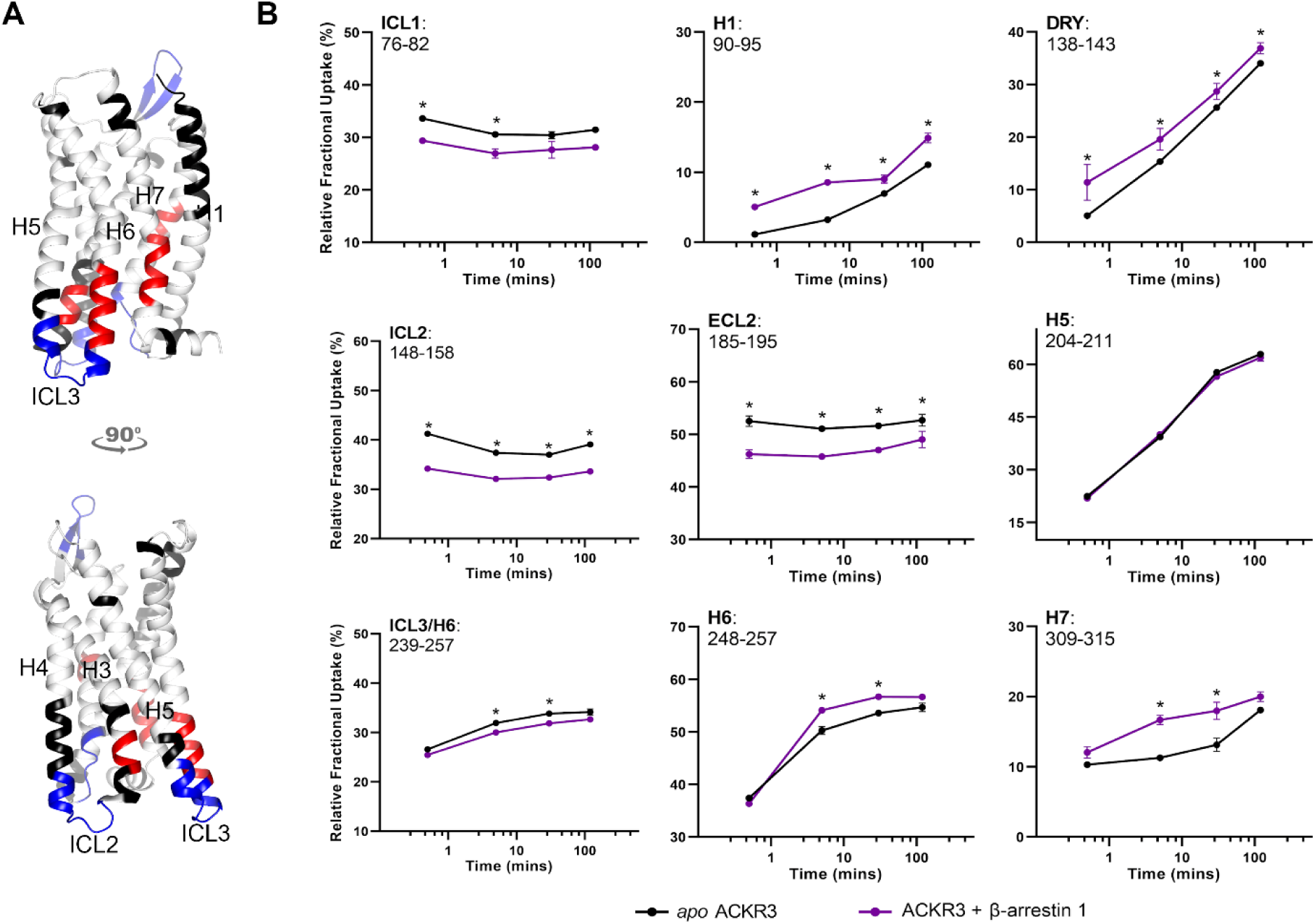
β-Arrestin 1 binding to ACKR3. **A)** Schematic representation of the protected (blue) and deprotected (red) regions of ACKR1 in the presence of β-arrestin 1 mapped onto the AF model of ACKR3 (for clarity, the N- and C- termini are not shown). This depicts reproducible and statistically significant ΔHDX in the presence of β-arrestin 1. Black regions represent regions with no sequence coverage. **B)** Associated deuterium uptake plots showing the relative uptake for peptides from *apo* or arrestin-bound ACKR3 across several deuteration time points and are representative of the region as indicated at the top of the plot. Statistically significant changes are represented by black stars and were determined using Deuteros 2.0 software^48^ (p ≤ 0.01). Uptake plots are the average and SD of 3 technical replicates from the same biological preparation of ACKR3.

The overall ΔHDX profile hints at a core engagement with β-arrestin 1, possibly in a similar fashion than those reported for the V2R^60^ and the NTSR1^61^, which both engage β-arrestin with all three ICLs. Since we do not have any available structures of ACKR3/β-arrestin complex, we generated AF models of this latter (Fig. S6). The 10 top-ranked AF models all showed core engagement with β-arrestin 1, where the middle loop and finger loop interact with the receptor intracellular pocket, ICL1, 2 and 3. Interestingly, comparison of these models with the ACKR3 apo conformation revealed an inward motion of the NPxxY motif in the β-arrestin-bound model, consistent with its decreased deuterium uptake. In addition, the multiple interactions observed between ACKR3 ICls and β-arrestin 1 correlate with the observed decrease in deuterium uptake for these loops in the presence of β-arrestin 1. Often ICL3 is not well resolved in high-resolution GPCR structures. However, ICL1 and ICL2 have been highlighted as important interfaces for β-arrestin binding. In the majority of published cryo-EM complexes, ICL2 sits in a defined cleft between the N and C domains of β-arrestin^60, 62–64^. The only known exception is NTSR1 where ICL1 takes this position instead^65^. Additionally, ICL1 has been shown to contribute to β-arrestin binding for all GPCR/β-arrestin complexes apart from that of the M2R^64^. The increased deuterium uptake observed at ICL2 in response to the agonist and the protection of all ICLs in the presence of β-arrestin 1 therefore suggests that these regions mediate the interaction with β-arrestin. On the other hand, the deprotection observed at the NPxxY motif indicates that this region plays a critical role for β-arrestin 1 binding, and additional experiments are required to better explore the structural basis of this impact.

## Discussion

Being an intrinsically β-arrestin-biased chemokine receptor, ACKR3 is an interesting subject for the study of GPCR functional selectivity. Its promiscuity for non-chemokine ligands and selectivity for chemokines add to its intriguing character. Combining HDX-MS and MD analysis, we identified the inactive state of ACKR3, as well as typical and atypical conformational features of ACKR3 activation (Fig. 5). It has been suggested that ACKR3 may have an intrinsically “active-like” structure^7^, given its high basal activity. Our data show that ACKR3 adopts typical class A inactive-active conformational changes involving H6 opening. This is triggered by an overall twist of the 7TM bundle upon agonist binding, which propagates allosterically to the intracellular side in a loosely-coupled manner. The inverse agonists, by contrast, stabilised the inactive state of the 7TM bundle through additional interactions in a sub-pocket between H2 and H7. β-arrestin 1 binding led to HDX deprotection of H2 immediately below the H2-H7 sub-pocket, which is likely a hot spot for ligand bias, as it was reported in other class A GPCRs^28, 57^. The small-molecule ligands bind deeper than CXCL12 and occupy different sub-pockets, which may explain the promiscuity of ACKR3 for diverse ligands.

**Figure 5.**
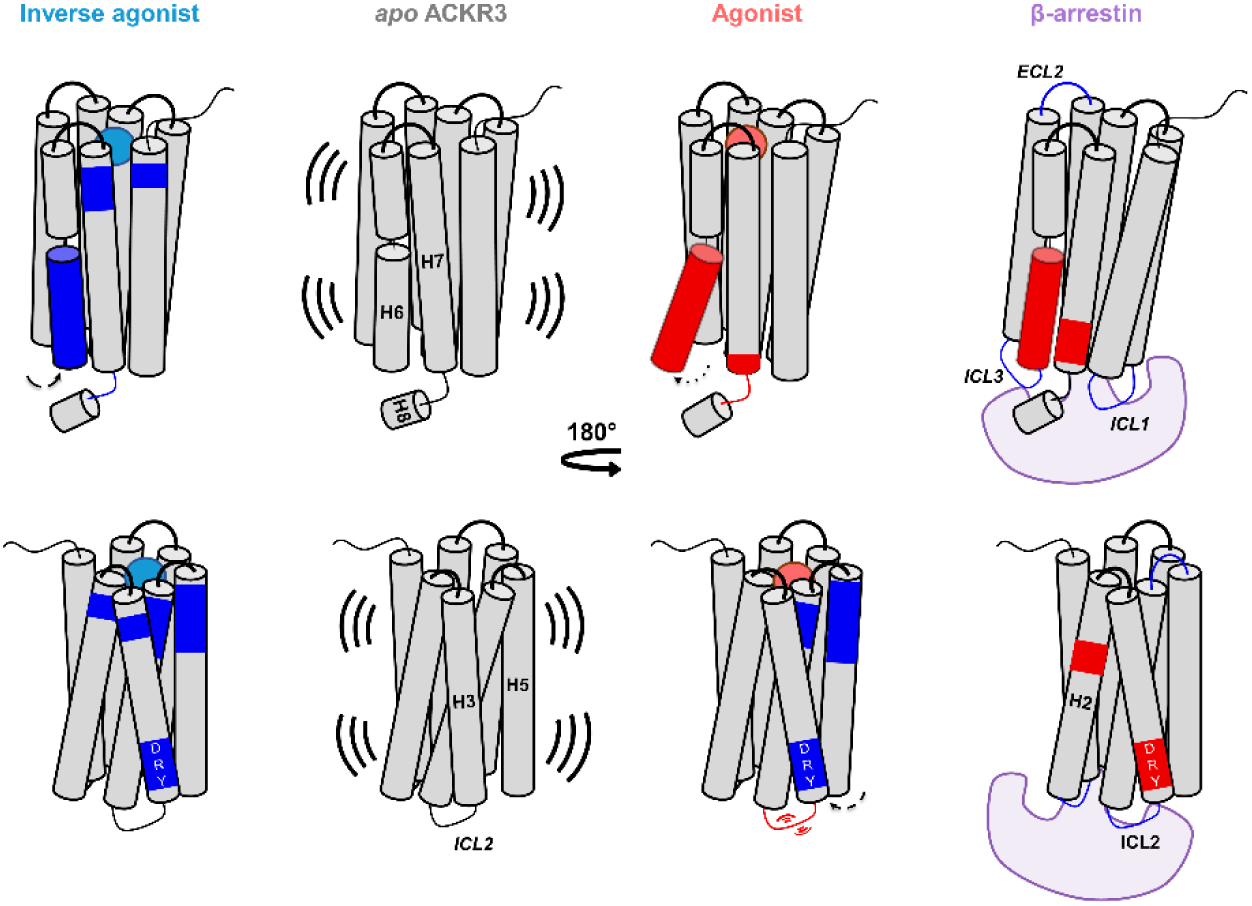
Scheme summarizing changes induced by binding of an agonist, inverse agonist and β-arrestin 1 to ACKR3. Activation through agonist binding induces an allosteric opening of H6 and resulted in increased flexibility and/or solvent exposure (red regions) of peptides spanning H6, H7 and ICL2. Inverse agonists resulted in decreased flexibility and/or solvent exposure (blue regions) through receptor constraint. β-arrestin 1 binding reduced the deuteration of all intracellular loops and resulted in increased deuteration of the NPxxY motif (H7), DRY motif (H3) and the conserved sodium binding site D90^2.50^ (H2). Arrows depict movements for the respective helices in response to different binders. Activation is associated with outward movement of TM6 and inward movement of H5 which contributes to protection of the DRY motif in the presence of the agonist.

ICL2 and the intracellular tip of H4 also exhibited significant deprotection upon agonist binding. As H6 moved outward, the contacts between H5-ICL3-H6 and H3 changed remarkably, leading to ICL2 unfolding. Since ICL2 forms a short α-helix in in most class A GPCRs and has been associated with G protein subtype selectivity of several receptors^36, 37, 54^, our data suggest that ICL2 unfolding might contribute to the lack of G protein signalling of ACKR3.

Another intriguing observation is that the DRY motif was protected upon ligand binding, even after opening of H6 in the presence of the agonist. It appeared to be shielded by Y232^5.58^, Y257^6.40^ and Y315^7.53^, which formed π-stacking in the active state. This feature is, to our knowledge, unique to ACKR3 in non-olfactory class A GPCRs. However, it is unclear whether the reduced accessibility of the DRY in ACKR3 contributes to its lack of G protein signalling. In the presence of β-arrestin 1, on the other hand, the DRY motif was slightly deprotected, suggesting that this motif is not shielded upon ACKR3 binding to β-arrestin 1.

In the presence of β-arrestin 1, all three ICLs of ACKR3 were unambiguously protected from deuterium exchange, suggesting therefore a core engagement with β-arrestin 1. Interestingly, peptides spanning the NPxxY motif as well as the intracellular face of H6 were deprotected upon β-arrestin 1 binding. Several studies have reported important roles of H7 and the NPxxY motif in class A GPCR signalling bias^28, 66–68^. Particularly, Y315^7.53^ is highly conserved^69^ and the Y315^7.53^A mutation almost completely abolishes β-arrestin recruitment by ACKR3^8^. The deprotection at the NPxxY motif therefore suggests that this region becomes more exposed or more flexible in the presence of β-arrestin. Further studies are required to better understand the functional significance of the NPxxY conformational change.

## Methods

### Expression and purification of ACKR3

For production in insect cells, the full-length gene of human ACKR3 was subcloned into pFastBac1 to enable infection of sf9 insect cells. The construct bore a hemagglutinin signal peptide followed by a Flag-tag preceding the receptor sequence. ACKR3 N13, N22 and N33 residues were substituted with a Glutamine in order to avoid N-glycosylation.

Flag-ACKR3 was expressed in sf9 insect cells using the pFastBac baculovirus system (Thermo Fisher Scientific). Cells were grown in suspension in EX-CELL 420 medium (Sigma-Aldrich) and infected at a density of 4 × 10^6^ cells/ml with the recombinant baculovirus. Flasks were shaken for 48 hours at 28°C, subsequently harvested by centrifugation (3,000*g*, 20 min) and stored at -80 until usage. Cell pellets were thawed and lysed by osmotic shock in a buffer containing 10mM Tris (pH 7.5), 1mM EDTA, 2 mg/ml iodoacetamide, 1μM ACKR3 agonist VUF11207 and protease inhibitors: 50μg/ml Leupeptin (Euromedex), 0.1mg/ml Bensamidine (Sigma-Aldrich) and 0.1mg/ml Phenylmethylsulfonyl fluoride (PMSF; Euromedex). Lysed cells were centrifuged for (38,400*g*, 10mins) and the resulting pellet was solubilised and dounce-homogenised 20x in buffer containing 50mM Tris (pH 7.5), 150mM NaCl, 2 mg/ml iodoacetamide, 1μM VUF11207, 0.5% (w/v) DDM (Anatrace), Cholesteryl hemisuccinate 0.1% (w/v) and protease inhibitors (50μg/ml Leupeptin, 0.1mg/ml Bensamidine and 0.1mg/ml PMSF). The homogenate was subsequently stirred for 1 h at 4 °C and centrifuged (38,400*g*, 30 min). The supernatant was then loaded onto M2 anti-Flag affinity resin (Sigma-Aldrich) using gravity flow. Resin was subsequently washed with 10 column volumes (CV) of DDM wash buffer containing 50mM Tris, 150mM NaCl, 0.1μM VUF11207, DDM 0.1% (w/v), Cholesteryl hemisuccinate 0.02% (w/v). Detergent was then gradually from DDM to LMNG (Anatrace) using increasing ratios of DDM wash buffer and buffer containing 50mM Tris, 150mM NaCl, 0.02μM VUF11207, 0.2% MNG (w/v), 0.05% CHS (w/v). Once detergent was fully exchanged, MNG and CHS concentration were steadily reduced to 0.005% and 0.001% respectively. ACKR3 was finally eluted in 50mM Tris, 150mM NaCl, 0.02μM VUF11207, 0.005% MNG (w/v), 0.001% CHS (w/v) and 0.4 mg/ml Flag peptide (Covalab). The eluate was concentrated using a 50kDa molecular weight cutoff (MWCO) concentrator (Millipore), then ACKR3 was purified by size exclusion chromatography (SEC) using a Superdex 200 Increase (10/300 GL column) connected to an ÄKTA purifier system (GE Healthcare) and eluted in buffer elution buffer without Flag peptide or VUF11207. Fractions containing monomeric ACKR3 were concentrated to between 20 and 25μM, aliquoted, flash-frozen and stored at -80°C prior to HDX experiments.

### Expression and purification of β-arrestin 1

A truncated form of β-arrestin1 was used in HDX experiments due to the higher basal activity of the protein. This construct was truncated at residue 382 and produced as follows. Competent BL21 *Escherichia coli* cells (Thermo Fisher Scientific) were transformed using a pET plasmid containing β-arrestin1 ΔC-ter containing a Twin-Strep tag at the N-terminus. Cells were cultured in LB with Kanamycin (37°C, 170 rpm) until an optical density of 0.6U was reached after which 0.025 mM Isopropyl-β-D-thiogalactopyranoside was added to induce cell expression. Cells were further incubated for 5hrs at 37°C and harvested by centrifugation (3000*g*). Pellets were stored at -80 prior to use. Protein purification was as follows. Cell pellets were thawed and resuspended in buffer containing Tris-HCl (pH8), 1mM EDTA, 200mM NaCl and 1mM β-mercaptoethanol and protease inhibitors Leupeptin (5μg/ml), benzamidine (10μg/ml) and PMSF (10μg/ml). Cells were then lysed by sonication then MgCl_2_ and benzonase were added to the lysate, 5mM and 2000 U respectively. Lysate was centrifuged (38,400*g*, 20mins, 4°C) and supernatant was supplemented with BioLock (0.75 ml/L) and loaded onto StrepTactin affinity resin at 4°C. The resin was then washed with a wash buffer containing 20mM Tris (pH8), 200mM NaCl and 100uM Tris(2-carboxyethyl)phosphine (TCEP). Wash buffer supplemented with 2.5mM desthiobiotin (IBA) was used to elute the protein. The eluate was then further purified by size exclusion chromatography using a Superdex 200 Increase (10/300 GL column) connected to an ÄKTA purifier system (GE Healthcare) in a buffer containing 20mM Hepes (pH 7.5), 200mM NaCl and 100μM TCEP. Eluted fractions containing purified β-arrestin were collected and concentrated using a 10kDa MWCO concentrator (Millipore). Aliquots were flash-frozen and kept at -80°C.

### HDX-MS experiments

HDX-MS experiments were performed using a Synapt G2-Si HDMS coupled to nanoAQUITY UPLC with HDX Automation technology (Waters Corporation). ACKR3 in LMNG detergent was concentrated up to 20-25uM µM and optimization of the sequence coverage was performed on undeuterated controls. Various quench times and conditions were tested, i.e. in the presence or absence of different denaturing or reducing reagents with or without longer trapping times to wash them out. The best sequence coverage and redundancy for ACKR3 were systematically obtained without the addition of any denaturing agents in the quench buffer. Mixtures of receptor: ligands were pre-incubated together to reach equilibrium prior to HDX-MS analysis. Analysis of freshly prepared ACKR3 apo, ACKR3: small ligands (1: 10 ratio) and ACKR3: β-arrestin (1: 1.2 ratio) mixtures were performed as follows: 3 µL of sample are diluted in 57 µL of undeuterated for the reference or deuterated last wash SEC buffer. The final percentage of deuterium in the deuterated buffer was 95%. Deuteration was performed at 20°C for 0.5, 2, 5, 30 and 120mins. Next, 50 µL of reaction sample are quenched in 50 µL of quench buffer (KH2PO4 50 mM, K2HPO4 50mM, 200 mM TCEP pH 2.3) at 0°C. 80 µL of quenched sample are loaded onto a 50 µL loop and injected on a Nepenthesin-2 column (Affipro) maintained at 15°C, with 0.2% formic acid at a flowrate of 100 µL/min. The peptides are then trapped at 0°C on a Vanguard column (ACQUITY UPLC BEH C18 VanGuard Pre-column, 130Å, 1.7 µm, 2.1 mm × 5 mm, Waters) for 3 min, before being loaded at 40 µL/min onto an Acquity UPLC column (ACQUITY UPLC BEH C18 Column, 1.7 µm, 1 mm × 100 mm, Waters) kept at 0°C. Peptides are subsequently eluted with a linear gradient (0.2% formic acid in acetonitrile solvent at 5% up to 35% during the first 6 min, then up to 40% and 95% over 1 min each) and ionized directly by electrospray on a Synapt G2-Si mass spectrometer (Waters). HDMSE data were obtained by 20-30 V trap collision energy ramp. Lock mass accuracy correction was made using a mixture of leucine enkephalin and GFP. For every tested condition we analysed two to three biological replicates, and deuteration timepoints were performed in triplicates for each condition.

Peptide identification was performed from undeuterated data using ProteinLynx global Server (PLGS, version 3.0.3, Waters). Peptides are filtered by DynamX (version 3.0, Waters) using the following parameters: minimum intensity of 1000, minimum product per amino acid of 0.2, maximum error for threshold of 10 ppm. All peptides were manually checked, and data was curated using DynamX. Back exchange was not corrected since we are measuring differential HDX and not absolute one. Statistical analysis of all ΔHDX data was performed using Deuteros 2.0^48^ and only peptides with a 99% confidence interval were considered.

### AlphaFold predictions

A SBGrid consortium installation of AlphaFold version 2.3^70, 71^ was run on a local server equipped with a NVIDIA Tesla P100 GPU in order to predict the structure of the ACKR3 monomer and ACKR3/β-arrestin1 heterodimer. The full databases were used, with max_template_date = 2023-04-09. The model_preset option was set to “monomer” for ACKR3 and to “multimer” for the heterodimer. All other parameters were left to their default values.

### MD simulations

Ligands were docked into ACKR3 models using Autodock Vina^72^. Four isomers per ligand were tested, including 2 diastereoisomers and 2 amine enantiomers. Pocket residues and ligand rotatable bonds were set as flexible. Top-ranked binding poses underwent REST2 MD simulations until a stable binding pose was obtained.

CHARMM-GUI was used to assign the side-chain protonation states and embed the models in a bilayer of POPC and cholesterol in 3:1 ratio. The systems were solvated in a periodic box of explicit water and neutralized with 0.15 M of Na^+^ and Cl^-^ ions. We used the Amber ff14SB^73^ GAFF^74^ and lipid21^75^ force fields, the TIP3P water model and the Joung-Cheatham ion parameters^76^. Effective point charges of the ligands were obtained by RESP fitting of the electrostatic potentials calculated with the HF/6-31G* basis set using Gaussian 16^77^.

After energy minimization, all-atom MD simulations were carried out using Gromacs2020^78^ patched with PLUMED2.3^79^. The LINCS algorithm^80^ was applied to constrain bonds involving hydrogen atoms, allowing for a time step of 2 fs. Each system was gradually heated to 310 K and pre-equilibrated during 10 ns of brute-force MD in the *NPT-*ensemble. The replica exchange with solute scaling (REST2)^46^ technique was used to enhance the conformational sampling. A total of 96 replicas of simulations were performed in the *NVT* ensemble. REST2 is a type of Hamiltonian replica exchange simulation scheme. Besides the original simulation, many replicas of the same system were simulated simultaneously. The additional replicas have modified free energy surfaces, in which the energy barriers are easier to cross than in the original simulation system. By frequently swapping the replicas and the original system during the MD, the simulations “travel” on different free energy surfaces and easily visit various conformational zones. Finally, only the samples on the original free energy surface are collected. The additional replicas are artificial to ease barrier crossing, which are discarded after the simulations. REST2, in particular, modifies the free energy surfaces by scaling (reducing) the force constants of the “solute” molecules in the simulation system. In this case, the protein was considered as “solute”–the force constants of its van der Waals, electrostatic and dihedral terms were subject to scaling in order to facilitate the conformational changes. The scaling factors were generated using the Patriksson-van der Spoel approach^81^ and effective temperatures ranging from 310 K to 1800 K. Exchange between replicas was attempted every 1000 simulation steps. This setup resulted in an average exchange probability of ∼30%. The simulation trajectories on the original unmodified free energy surface were reassembled and analysed.

Several independent REST2 MD runs were performed on different docked poses of each ligand until a stable binding pose was obtained, in which the ligand RMSD was < 2 Å during at least 30 ns. Discarding the unstable binding poses, the remaining simulations were prolonged until the HDX profile could be reproduced. A total of > 1.5 milliseconds of REST2 MD simulations (including replicas) were performed. The final production runs contained 60 ns - 100 ns × 96 replicas for each system. The last 30 ns were used for HDX logPF calculations using HDXer^47^. All the atoms, with the exception of water molecules, were taken into account. The remaining parameters in HDXer were set to their default values. Principal component analysis was performed with bio3d^82^, after an alignment on the most invariant region of the receptor. The most invariant region was identified by iterations of structural superposition, excluding the most variable regions after each round.

### Analysis of ACKR3 ligand binding site by site-directed mutagenesis

The binding sites of the agonist VUF15485 and the inverse agonist VUF16840 were also probed by site-directed mutagenesis. N-terminally HA-tagged ACKR3 mutants, D179^4.60^N, E213^5.39^Q, D275^6.58^N, Q301^7.39^E and Q301^7.39^A were generated by PCR-based mutagenesis, sequence-verified and expressed in HEK293T cells, as described previously^42^. The binding of VUF15485 and VUF16840 to wild-type and mutant ACKR3 was measured in [^3^H]VUF15485 (in-house synthesised) radioligand binding studies^42^. Briefly, membranes expressing the wild type or mutant HA-ACKR3 were incubated with [^3^H]VUF15485 and increasing concentrations VUF16840 or VUF15485. All dilutions were prepared in HBSS supplemented with 0.2% BSA. Membranes and radioligands were optimized to measure specific binding of [^3^H]VUF15485 to the ACKR3 and mutants, with 1000 - 5000 dpm. Membrane protein contents ranged between 4 μg and 80 μg per well and [^3^H]VUF15485 ranged between 2 nM and 20 nM. All conditions were performed in triplicate on a 96-well plate. Binding reactions were incubated for 2 hours and then terminated by washing the solutions over a PEI-coated GF/C filter using a cell harvester (Perkin Elmer) followed by wash steps using ice-cold buffer [50 mM HEPES, 1.2 mM CaCl_2_, 5 mM MgCl_2_, 0.5 M NaCl, pH 7.4 at 4°C]. Filter-bound [^3^H]VUF15485 was measured by adding 25 µL Microscint-O per well to the dried GF/C-plate and radioactivity was consecutively quantified using the Wallac Microbeta counter (Perkin Elmer). The sigmoidal dose-dependent displacement curves of [^3^H]VUF15485 binding by unlabelled ligands were analysed using Graphpad Prism 8, by fitting the data to a one-site binding model to determine the pIC_50_. Binding affinities (K_i_ values) for VUF15485 and VUF16840 were determined using the Cheng-Prusoff equation^83^.

## Acknowledgments

This research was funded by European Union’s Horizon2020 Marie Skłodowska-Curie Actions (MSCA) to OO, RL, TD, SG and CB who are part of the Marie Skłodowska-Curie Innovative Training Network ONCORNET2.0 “ONCOgenic Receptor Network of Excellence and Training” (MSCA-ITN-2020-ETN). Technical assistance of Drs. Simon Mobach is highly valued. The work of MA-S, RL and CdG was supported by European Union’s Horizon2020 MSCA Program under grant agreement 641833 [ONCORNET]. Mass spectrometry experiments were carried out using the facilities of the Montpellier Proteomics Platform (PPM, BioCampus Montpellier). This work was supported by the regional funds FEDER/Région Occitanie, MUSE, Labex EpiGenMed and the French Agence Nationale de la Recherche (project ANR-21-CE44-0007-01, LEUKOCEPTOR).

## Author contributions

OO expressed and purified the proteins under the supervision of SG. OO, EDN and TG collected and analysed the HDX-MS data under the supervision of CB. CA implemented and adjusted the HDXer python package under the supervision of XC. XC performed and analysed ligand docking and MD simulations. CL performed the AF modelling. MA-S, RB, BZ and CdG performed the initial ligand docking and site-directed mutagenesis experiments under the supervision of RL. TD, SG and CB initiated the project. CB planned and supervised the research. OO, XC and CB wrote the manuscript with input from all co-authors.

