## Supplementary Material for "Conformational dynamics underlying Atypical Chemokine Receptor 3 activation"

**Figure S1**: Relative fractional uptake for apo ACKR3 peptides.

**Figure S2**: Movements of H6 intracellular tip during the MD simulations initiated from the AF model.

**Figure S3**: Coverage maps and representative heat maps for tested ligands.

**Figure S4**: Protection of the DRY motif in VUF15485-bound ACKR3 during MD simulations.

**Figure S5**: Principal component analysis (PCA) of the MD trajectories.

**Figure S6**: Alphafold models of the ACKR3 – β-arrestin 1 heterodimer in comparison with the ACKR3 monomer model.

**Table S1**: Binding affinity of VUF15485 and VUF16840 for ACKR3 and selected binding site mutants receptors as measured by [$^3$H]VUF15485 binding.

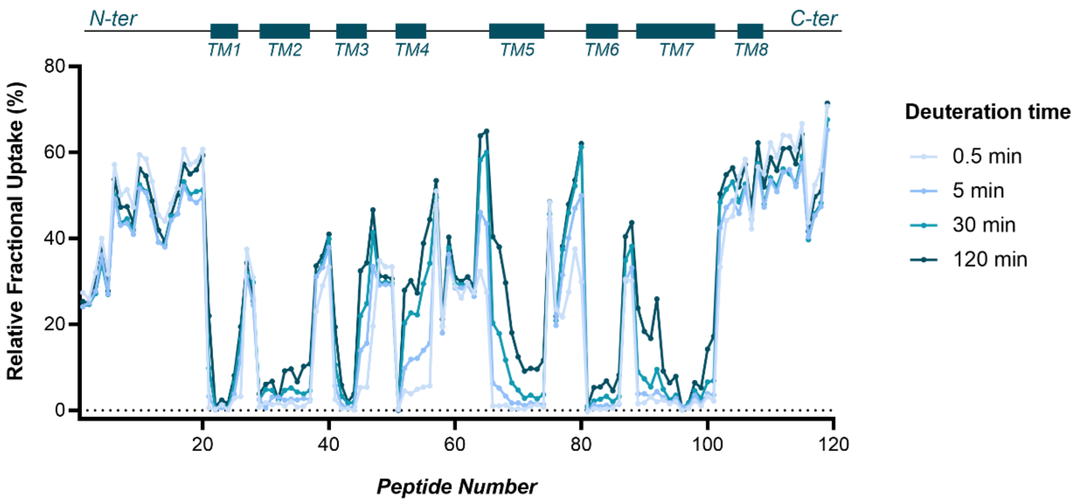


**Supplementary figure 1: Relative fractional uptake for apo ACKR3 peptides.**

Plot of representative peptides identified in HDX experiments and their total % relative fractional uptake across tested deuteration times (0.5, 5, 30 and 120 min). Graph shows uptake for apo-ACKR3, detected peptide regions are according to the included scheme.

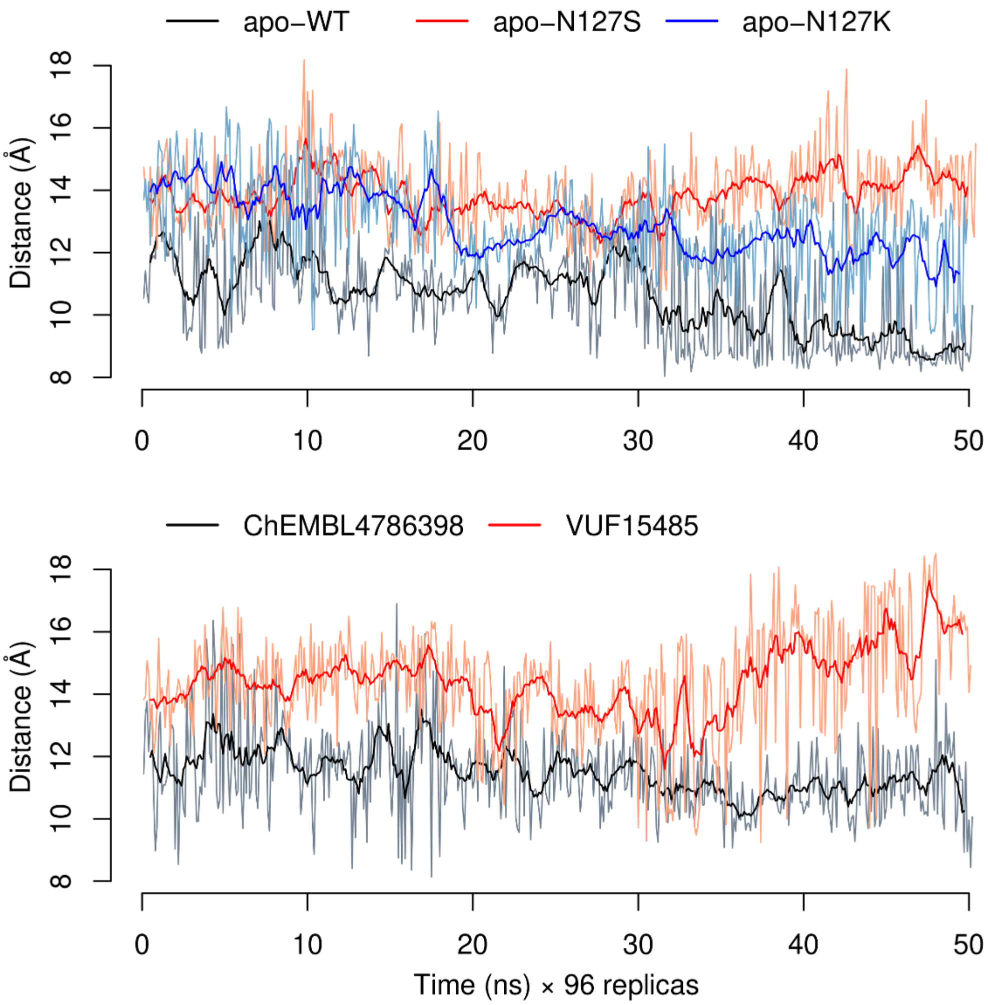


**Supplementary figure 2: Movements of H6 intracellular tip during the MD simulations initiated from the AF model.**

Plots of the R142$^{3.50}$-E246$^{6.29}$ Cα distance in each system.

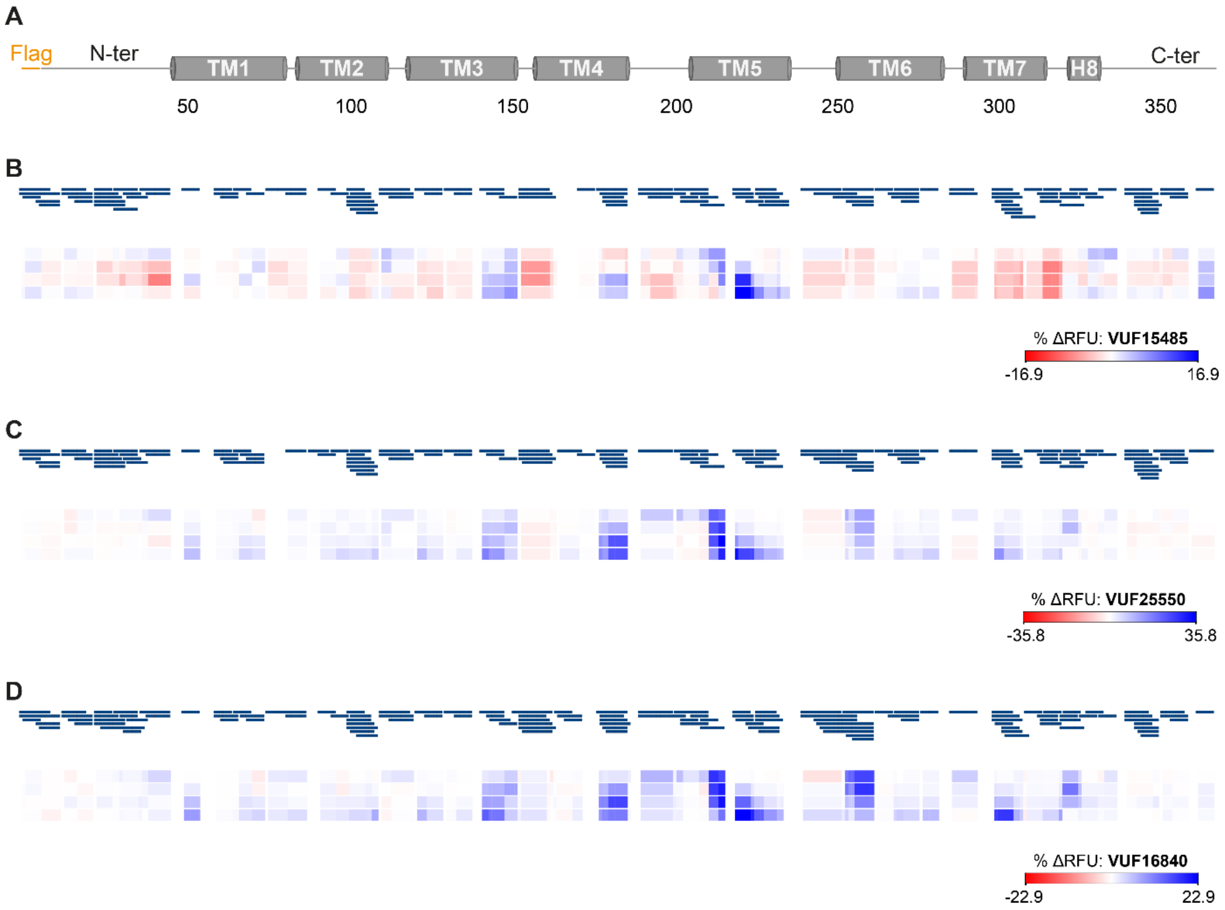


**Supplementary figure 3: Coverage maps and representative heat maps for tested ligands.**

**A)** Scheme representing transmembrane helices in correspondence to residue numbers. **B-D)** Representative relative fractional uptake differences (%) and coverage maps for HDX analysis of ACKR3 in the presence of all tested ligands (apo – ligand-bound ACKR3). Deuteration was performed at 0.5, 5, 30 and 120 mins for all states. For additional information, see supported Data S1 and S2 (xls files).

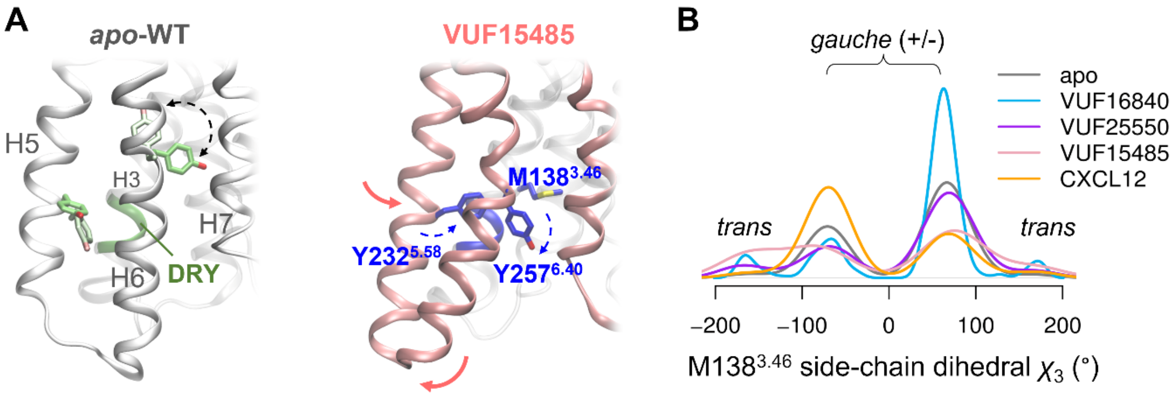


**Supplementary figure 4: Protection of the DRY motif in VUF15485-bound ACKR3 during MD simulations.**

**A)** MD simulations of ACKR3 in it *apo* and VUF15485-bound forms, showing that the DRY motif is shielded from the solvent by Y232$^{5.58}$ and Y257$^{6.40}$ upon agonist binding despite the H6 opening. While H6 moved outward, H5 moved inward and provided additional protection to the DRY motif through intramolecular contacts. In the *apo* form, Y232$^{5.58}$ pointed outside the 7TM bundle, while Y257$^{6.40}$ pointed upward or laid above the DRY motif. In each form, Y257$^{6.40}$ showed two orientations during the simulations. **B)** Density distribution of M138$^{3.46}$ side-chain dihedral angle $\chi_3$ during the MD simulations.

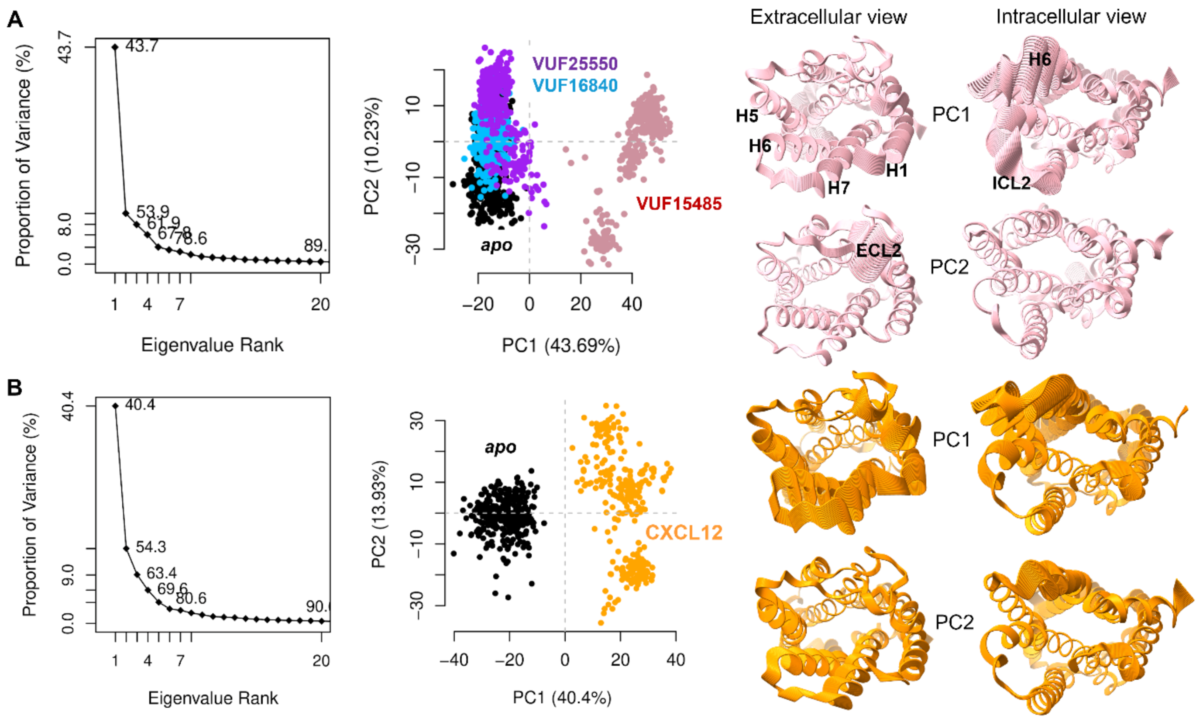


**Supplementary figure 5: Principal component analysis (PCA) of the MD trajectories.**

**A)** Grouped PCA of 4 MD trajectories (*apo* ACKR3 and bound with ligands), which allows direct comparison of the movements in the 4 systems. Principal component (PC) 1 corresponds to VUF15484-induced 7TM twist around the pocket and the receptor conformational changes on the intracellular side. PC1 corresponds to ECL2 movements in the systems, which may be artefacts due to the truncated N-terminus in the MD simulations. **B)** Grouped PCA of *apo* ACKR3 *versus* the CXCL12-bound form. Both PC1 and PC2 both exhibit CXCL2-induced 7TM twist and intracellular conformational changes, similar to the PC1 in (A). PCA was performed on the Cα atoms of the 7TM domain.

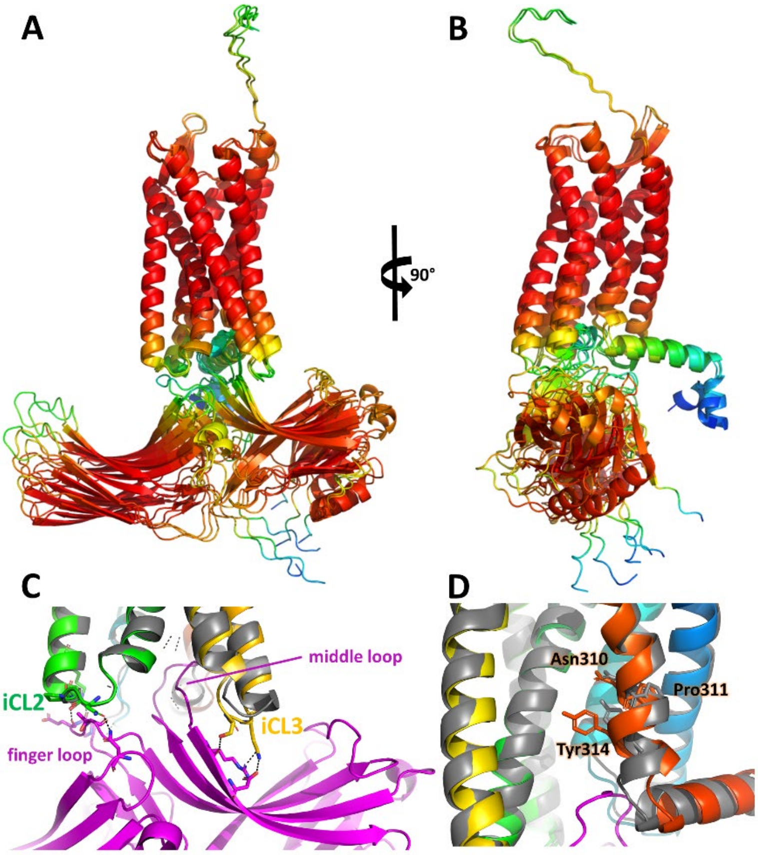


**Supplementary figure 6: Alphafold models of the ACKR3 – β-arrestin 1 heterodimer in comparison with the ACKR3 monomer model**.

**A)** and **B)** the top 3 AF models in 2 views rotated by 90°. Models were superimposed by alignment on the ACKR3 chain, and are shown in cartoon representation and coloured by AF predicted local distance difference test (pLDDT) score. **C)** Close up view of the interaction interface between ACKR3 and β-arrestin 1, highlighting the polar contacts formed between β-arrestin 1 and ACKR3 ICL2 and 3. The β-arrestin 1 is coloured in magenta while ACKR3 is coloured from blue (N-terminus) to red (C-terminus). The ACKR3 monomeric model is superimposed onto the model of the complex and coloured in grey. **D)** Superimposed ACKR3– β-arrestin 1 and ACKR3 models, highlighting the motion of the NPxxY motif. The colour code is the same as in C.

**Table S1: Binding affinity of VUF15485 and VUF16840 for ACKR3 and selected binding site mutants receptors as measured by [$^3$H]VUF15485 binding.** Binding affinities ($K_i$) were determined from $pIC_{50}$ using the Cheng-Prusoff equation. All values depict the mean ± SEM of (N) experiments.

| **$pK_i$ ± SEM (N)** | **WT** | **D179N$^{4.60}$** | **E213Q$^{5.39}$** | **D275N$^{6.58}$** | **Q301E$^{7.39}$** | **Q301A$^{7.39}$** |
|---|---|---|---|---|---|---|
| **VUF15485** | 7.9 ± 0.1 (18) | 6.4 ± 0.2 (7)** | 7.9 ± 0.1 (7) | 8.1 ± 0.2 (4) | 7.5 ± 0.2 (4) | 8.0 ± 0.2 (8) |
| **VUF16840** | 9.3 ± 0.0 (13) | 7.5 ± 0.2 (7)** | 8.3 ± 0.2 (7)** | 9.2 ± 0.1 (4) | 8.7 ± 0.3 (4)* | 8.7 ± 0.1 (8)* |

* P < 0.05 compared to WT
** P < 0.001 compared to WT
